# Quantification and Diagnostic Relevance of Blood and Heme-Mediated Inhibition of Prion Detection by RT-QuIC

**DOI:** 10.1101/2025.04.21.649906

**Authors:** Robert B. Piel, David A. Schneider

## Abstract

Prion diseases are characterized by misfolding of prion protein (PrP) from correctly folded PrP^C^ to a disease-associated form, PrP^D^. Real-time quaking-induced conversion (RT-QuIC), detects prions by “seeding” reaction mixtures, which contain PrP^C^, with samples suspected to contain prions, resulting in PrP^D^ amplification. The assay is sensitive to inhibition by tissue constituents, including blood. Heme, a cofactor of hemoglobin (Hb), has been shown to bind PrP in an isoform-specific manner and to affect the stability of other pathogenic amyloids. In the present study, tissue samples from scrapie-positive sheep were used to seed RT-QuIC reactions in the presence of heme—as free hemin, as a cofactor of Hb, and as present in whole blood. At equivalent heme concentrations, the inhibitory action of free heme was the least and that of blood the greatest, suggesting other components of Hb and whole blood have additional inhibitory actions. We also demonstrate that this inhibition of RT-QuIC acts through disruption of the recombinant PrP^C^ assay substrate, rather than destruction of PrP^D^ seeds. Lastly, heme concentrations were measured in several ruminant tissues. Heme levels exceeded inhibitory thresholds in nearly all types of intact tissue but were reduced below inhibitory levels at a 1:1000 dilution of most tissue types, with whole blood being one of a few notable exceptions. Our results suggest that detection of PrP^D^ seeding activity is not precluded by exposure to heme in tissue samples, but that the final heme concentration introduced into the RT-QuIC assay mixture is the critical factor that impacts detection sensitivity.

## Introduction

In prion diseases, which include scrapie in sheep and goats, chronic wasting disease (CWD) in cervids, and various others in both agricultural animals and humans, initial transmission is followed by a prolonged asymptomatic incubation period which can often last years. As infectious prions are shed during preclinical infection(1, 2), these animals represent a critical management challenge to efforts targeting highly transmissible prion diseases. These include the final stages of eradication for classical scrapie in the United States, as well as responses to the continuing expansion of CWD across North America. Currently, the diagnosis of prion disease in small ruminants and cervid species is accomplished by immunoassay using lymphoid tissues typically collected postmortem. A reliable and highly sensitive assay for early-stage prion disease using peripheral tissues or blood would greatly aid in disease management efforts.

A defining feature of prion infection is the replication of the infectious unit (the prion) by templated misfolding of natively expressed prion protein (PrP) from its normal “cellular” form (PrP^C^) to a misfolded “disease” form (PrP^D^)(3). Amplification assays such as RT-QuIC exploit this inherent mechanism of replication to enable detection of miniscule amounts of misfolded protein that may otherwise elude conventional methods such as immunoassay. In RT-QuIC, enhanced detection is accomplished by exposing a reaction mixture containing correctly folded recombinant prion protein substrate (rPrP^C^) to test samples which may contain prions(4, 5). In the case of a positive sample, PrP^D^ in the sample acts as a template or “seed” for the conversion of the rPrP^C^ substrate to the misfolded form. The assay then proceeds in cycles of shaking, through which the newly misfolded substrate can also participate in seeding further conversion, thereby amplifying the amount of misfolded prion protein in the reaction mixture to detectable levels.

The RT-QuIC assay has shown exceptional sensitivity for the detection of prions, where seeding activity can be detected in positive samples that have been diluted more than ten billion-fold from the starting tissue(5). However, a significant hurdle facing the assay is that RT-QuIC is inhibited by various constituents naturally present in tissues and bodily fluids. A common practice has been to dilute tissue samples 1000-fold (10^−3^) before use in the assay(5–7). While this technique does succeed in circumventing a large part of tissue-mediated inhibition, it also inherently limits the potential sensitivity of the assay, thereby limiting the potential of amplification assays to enhance the detection of early infections.

In addition to “tissue” generally, constituents of blood and, in particular, red blood cells, are known to strongly inhibit the RT-QuIC assay (8–11). This is problematic both because blood is a near ubiquitous contaminant of most tissues, and because blood itself is known to harbor infectious prions (12–16) and is thus a very attractive sample type for live animal testing that could be repeated over time. While some progress has been made in detecting bloodborne prions by RT-QuIC with pre-analytic processing techniques such as selective precipitation(12) and affinity purification(10), it is clear that some level of inhibition is still present relative to non-blood-exposed samples(11). In contrast to other cell types, red blood cells, or erythrocytes, are unique in that >95% of their dry mass is made up of hemoglobin (Hb)(17, 18). Critical to its oxygen carrying function, each tetrameric Hb protein contains four heme molecules as cofactors. Free heme (hemin in its oxidized form) is a highly reactive porphyrin molecule which is known to cause oxidative damage(19, 20) when its normal cellular niche is disrupted, such as after hemolysis or ischemia reperfusion injury, conditions that are variably present during tissue sampling, storage, and homogenization. In addition to its potential to cause oxidative damage, heme also binds to PrP in an isoform-sensitive manner(21–23). While physiologic functions of PrP^C^-heme binding have been proposed for (21, 22), the implications for misfolding assays are not entirely clear. In addition to erythrocyte mediated inhibition of RT-QuIC(8), hemin itself has previously been shown to inhibit the detection of prions by another type of misfolding assay, the protein misfolding cyclic amplification (PMCA) assay (24). Furthermore, hemin has also been shown to influence the structures and stability of other disease-associated amyloids, including amyloid-β of Alzheimer’s disease(25, 26), α-synuclein of Parkinson’s disease(27), and lysozyme of lysozyme amyloidosis(28).

The present study aims to elucidate the extent to, and processes by which, blood and its major components, hemoglobin and heme, inhibit the RT-QuIC assay in the sensitive-detection of prions in tissue samples. This knowledge may ultimately allow for mitigation of this inhibition and the more sensitive detection of prions in blood-containing samples and/or blood directly.

## Methods

### Tissue preparation

The tissue samples used in this study are from a frozen archive of tissues collected from naturally infected and experimentally infected small ruminants. All animals were acquired, housed, and use procedures approved by the Institutional Animal Care and Use Committee of Washington State University (Animal Subjects Approval Forms: 4575, 6665, 6689, 6692). All tissue samples were collected after humane euthanasia and were same-day processed and frozen (−80 ℃) until use.

Sheep brain homogenates for RT-QuIC seeding (#4789 (pos), #4799 (neg)) were prepared as 10% (w/v) homogenates in 1× phosphate-buffered saline (PBS) using a rotor stator homogenizer (GLH-01, Omni International) with single use plastic probe tips.

Blood samples for RT-QuIC inhibition and rPrP^C^ binding studies were collected from a scrapie-naïve sheep (#5047) in 10 ml EDTA Vacutainer tubes (Becton Dickinson). Tissue samples for heme quantification were collected from two sheep (#4645, #4649) and one goat (#5010G). Tissues collected included brainstem, cerebellum, tonsil, retropharyngeal lymph node (RPLN), spleen, kidney, liver, diaphragm, skeletal muscle, rectal mucosa, cerebrospinal fluid (CSF), and whole blood collected with ACD anticoagulant (8:60 ml ACD Formula A; Fenwal). Sheep placental cotyledons were also collected, with 5 individual cotyledons gathered from each of 10 placentas (#1266-1275). Tissues for heme quantification experiments, excluding blood and CSF, were prepared as 10% (w/v) homogenates in 1× PBS using 0.7 mm Zirconia beads (BioSpec 11079107zx) in a bead beating grinder (Fast Prep 24 – MP bio). CSF was prepared by centrifugation at 500 x g for 10 min and the resulting supernatant collected. Tissue homogenate, CSF supernatant, and whole blood stocks were stored at −80 °C, with working sub-aliquots stored short term at −20 °C.

Homogenate concentrations in this manuscript are described as dilutions relative to intact tissues, i.e., a 10% (w/v) homogenate is represented as a 10^−1^ dilution.

### Heme and Hb solution preparation

Hemin (Hemin-Cl; EMD Millipore) and Hb (Hemoglobin from bovine blood; Sigma Aldrich) stock solutions were prepared gravimetrically from dry reagents.

Hemin was solubilized in 0.1 M NaOH and subsequently diluted at least 100-fold in seed dilution (SD) buffer [1× PBS pH 7.4 + 0.1% SDS + 1× N-2 media supplement (Gibco-Fisher)] to amend pH; hemin was found to be soluble in this buffer to at least 200 µM.

Hb was prepared in 1× PBS pH 7.4 for quantification standards or in SD buffer for RT-QuIC spiking. For Hb, concentrations are calculated and reported as monomeric Hb.

ApoHb was prepared from Hb stock solutions by acid-acetone extraction(29, 30). A solution of 20 mM Hb was added dropwise with constant stirring to approximately 30 volumes of acidified acetone (0.2% v/v 12N HCl) at −20°C. The resulting precipitate was collected by centrifugation at 1000 × g for 15 min and resuspended in ddH_2_O. The resulting solution was then successively dialyzed against ddH_2_O, 1.6 mM sodium bicarbonate, and 1x PBS pH 7.4. Following dialysis, residual precipitate was removed by centrifugation at 3000 x g for 10 min. ApoHb concentrations were estimated by UV-vis absorbance at 280 nm using ε_280nm_ = 0.0162 M^−1^ (29, 31). ApoHb dilutions prepared using this extinction coefficient were also compared to (w/v) holoHb standards by SDS-PAGE and Coomassie staining.

### Heme quantification in blood and tissues

Heme concentrations in whole blood were measured by alkaline detergent hematin (ADH) assay(32, 33). Dilutions of whole blood and Hb standards were prepared in 1× PBS. 20 ul of each were then added to 180 µl of a buffer consisting of 0.1 M NaOH + 2.5 % w/v Triton-X100. UV-vis absorbance spectra were recorded and blood-heme concentrations calculated by comparison to the Hb standard curve at 575 nm.

Heme concentrations in sheep and goat tissues were measured by oxalic acid fluorescence assay(34, 35). Dilutions of tissue homogenates and Hb standards were prepared in 1× PBS. 20 µl of each were then added to 980 µl of 2 M oxalic acid. The sample preparations were then split and 500 µl was incubated at 100 °C for 30 min while the other half was maintained at room temperature. Samples were then measured for fluorescence using an excitation wavelength of 400 nm and emission of 662 nm. For each sample, the room temperature measurements were subtracted from the 100 °C incubated measurements to control for any non-heme-derived fluorescence present in the samples. Measurements from the Hb standard curve were then used to calculate the concentration of heme for tissue samples.

UV-vis and fluorescence measurements were performed using a CLARIOstar microplate reader (BMG Labtech).

### RT-QuIC

RT-QuIC was performed using hamster-sheep chimeric rPrP^C^ substrate (Syrian hamster residues 23 to 137 [accession no. K02234] followed by sheep (R154, Q171) residues 141 to 234 [accession no. AJ567988]). Protein was expressed in DE3 Escherichia coli using the pET41 vector and Overnight Express Autoinduction System 1 (Novagen, Madison, WI). rPrP^C^ was purified from inclusion bodies as described by Orrú et al. (4). Briefly, inclusion bodies were solubilized in guanidine hydrochloride, purified using nickel immobilized metal affinity chromatography (IMAC), and refolded on the resin with a gradient of guanidine hydrochloride using an AKTA Pure FPLC (Cytivia). Following elution by imidazole gradient, rPrP^C^ was dialyzed into 10 mM Na_2_PO_4_ (pH 5.8) and stored at −80°C.

RT-QuIC reactions consisting of 2 μl seed material and 98 μl RT-QuIC assay buffer [10 mM NaPO_4_ pH 7.4, 300 mM NaCl, 1 mM EDTA, 10 μM ThT, 0.1 mg/mL rPrP^C^] per well were carried out at 42°C with alternating cycles of 1 min double orbital shaking at 700 rpm and 1 min rest for 100 h total using FLUOstar or CLARIOstar microplate readers (BMG Labtech). ThT fluorescence was measured using 20 flashes/well, bottom read, with an excitation wavelength of 450 ± 10 nm and emission wavelength of 480 ± 10 nm.

For RT-QuIC, seed homogenate dilutions were prepared in seed dilution (SD) buffer [1× PBS pH 7.4 + 0.1% SDS + 1× N-2 media supplement (Gibco-Fisher)]. In RT-QuIC reactions containing hemin, Hb, or blood, inhibitors were spiked into the seed homogenate dilutions such that the concentrations named represent the final concentrations present in the seed material. For blood, reported concentrations represent the heme/monomeric Hb concentration present in each dilution of whole blood.

### Methods used to evaluate RT-QuIC data

Each 100-hour record of ThT fluorescence measurements was exported to an excel spreadsheet. A custom script (Python version 3.10.9) was written to import, merge, and export as one comma-delimited data file all excel files relevant to a given experiment. Each experiment’s data file was then imported into SAS (SAS version 9.4) where various SAS procedures (PROCs) were applied to transform data, detect and characterize reactions, perform statistical analyses, and produce graphs for presentation.

RT-QuIC reaction data including excel exports of raw ThT fluorescence measurements and graphs depicting raw ThT fluorescence curves are available from the National Agricultural Library Ag Data Commons database (https://figshare.com/s/dbfd57004f51ffb7c64b). Examples of raw ThT fluorescence curves can be seen in Fig. 7a of this manuscript.

Given some extreme effects of the conditions of this study on the morphology and variability in the fluorescence data, custom algorithms were created and uniformly applied to each well’s data to detect reactions and evaluate morphologic features. The fluorescence readings were first regressed over time (PROC TRANSEG) to produce its penalized b-spline and upper 99.9% confidence limit. This resulted in the local reduction in variability and liberal upper confidence limit. Subsequently, a moving estimate of the fluorescence trend (ThT_trend_) was calculated as the leading 1-hour median of the penalized b-spline (PROC EXPAND). Similarly, a moving estimate of a critical threshold value (ThT_crit_) was calculated as the preceding 5-hour median of the fluorescence upper confidence limit offset by an additional 30-minutes. Thus, at a given time point, a relatively short forward-biased estimate of ThT_trend_ was compared to a local window estimate of the preceding baseline fluorescence (ThT_crit_). The lag time of a reaction was defined as the time at which ThT_trend_ first exceeded ThT_crit_. The reaction height at a given time point was defined as the baseline (ThT_crit_)-subtracted fluorescence.

Because of the inhibitory conditions being tested, reactions frequently could not be detected in all replicates. These instances were captured graphically by assigning them to a not-detected (ND) reference line placed at an arbitrary post-assay time of 110 hours. For statistical analysis, all times (lag and ND times) were first ranked (PROC RANK) and then analyzed using a generalized linear model based on the gamma distribution (PROC GLIMMIX). Occasional extreme outliers were identified and removed from analyses based on a panel of Studentized residual plots. All final models were well fit by the gamma distribution. Post-hoc analyses consisted of pre-planned comparisons of interest using the modeled least squares means, variation, and the Kenward-Roger (KR2) method of degrees freedom estimation for unbalanced data. The family-wise error rate of multiple comparisons was controlled using the adjustment method of Holm (i.e., stepdown Bonferroni). Significance was accepted at a *P* < 0.05.

### Heme-rPrP interaction

Hemin-rPrP interactions were quantified by UV-Vis spectral shift. RT-QuIC buffer without ThT [10 mM NaPO_4_ pH 7.4, 300 mM NaCl, 1 mM EDTA, 0.1 mg/ml rPrP^C^] was spiked with mock sample buffer (1X PBS + 0.1% SDS) or 1X PBS containing hemin at varying concentrations. Due to rPrP precipitation observed upon addition of the SDS+ buffer at 0 µM hemin, the 1× PBS buffer condition was used for binding ratio experiments. Matched hemin-only solutions were also prepared where hemin solutions were spiked into RT-QuIC buffer without ThT or rPrP^C^ [10 mM NaPO_4_ pH 7.4, 300 mM NaCl, 1 mM EDTA]. Differential spectra were taken, subtracting hemin-only spectra from hemin-rPrP^C^ spectra. Evolution of a differential peak at 416 nm and a valley at 385 nm was observed. The difference between these wavelengths was plotted against the hemin:rPrP^C^ molar ratio to show dose dependent evolution of the shift and eventual saturation of rPrP^C^ with hemin. Interaction of rPrP^C^ with Hb or Blood was tested by identical methods to investigate the possibility of heme transfer and/or binding to rPrP^C^ from either source.

### Seed exposure to Hb and Blood

10% brain homogenate from a scrapie-positive sheep was mixed 1:1 with whole blood, Hb solution, or PBS and incubated at 4 °C for 24 hours or 7 days. Prior to incubation, blood was lysed by sonication for 2 pulses of 30 seconds each in a water bath sonicator (Qsonica) at 180 W. Hb solution concentration was matched to that of whole blood as measured by ADH assay. Following incubation, mixtures were diluted in SD buffer and used as RT-QuIC seed material as described above.

### rPrP^C^ stability in RT-QuIC buffer

RT-QuIC assay buffer (−ThT) containing 0.1 mg/ml rPrP^C^ was incubated with mock seed samples containing hemin, Hb, or blood. UV/Vis spectra were then recorded on a CLARIOstar microplate reader (BMG Labtech). Samples were subsequently incubated for 24 h at 42 °C. Following incubation, samples were centrifuged at 21000 × g for 10 min and the supernatant was assayed for rPrP^C^ by SDS-PAGE and Coomassie staining.

## Results and Discussion

### Relative Inhibitory Effects of Blood, Hemoglobin, and Hemin on Prion RT-QuIC

To quantify the inhibitory effects of heme, Hb, and whole blood, a matrix of RT-QuIC assay conditions was assembled where scrapie positive sheep brain homogenates in dilutions ranging from 10^−3^ to 10^−6^ were spiked with heme, Hb, or whole blood at concentrations ranging from 0 to 200 µM (Fig 1). These concentrations represent the amounts of heme or monomeric Hb present in the seed material prior to RT-QuIC analysis, where 2 µl of seed material is then introduced to 98 µl of reaction buffer. For whole blood, the reported concentrations describe the amount of monomeric Hb the dilution contains.

**Figure 1.**
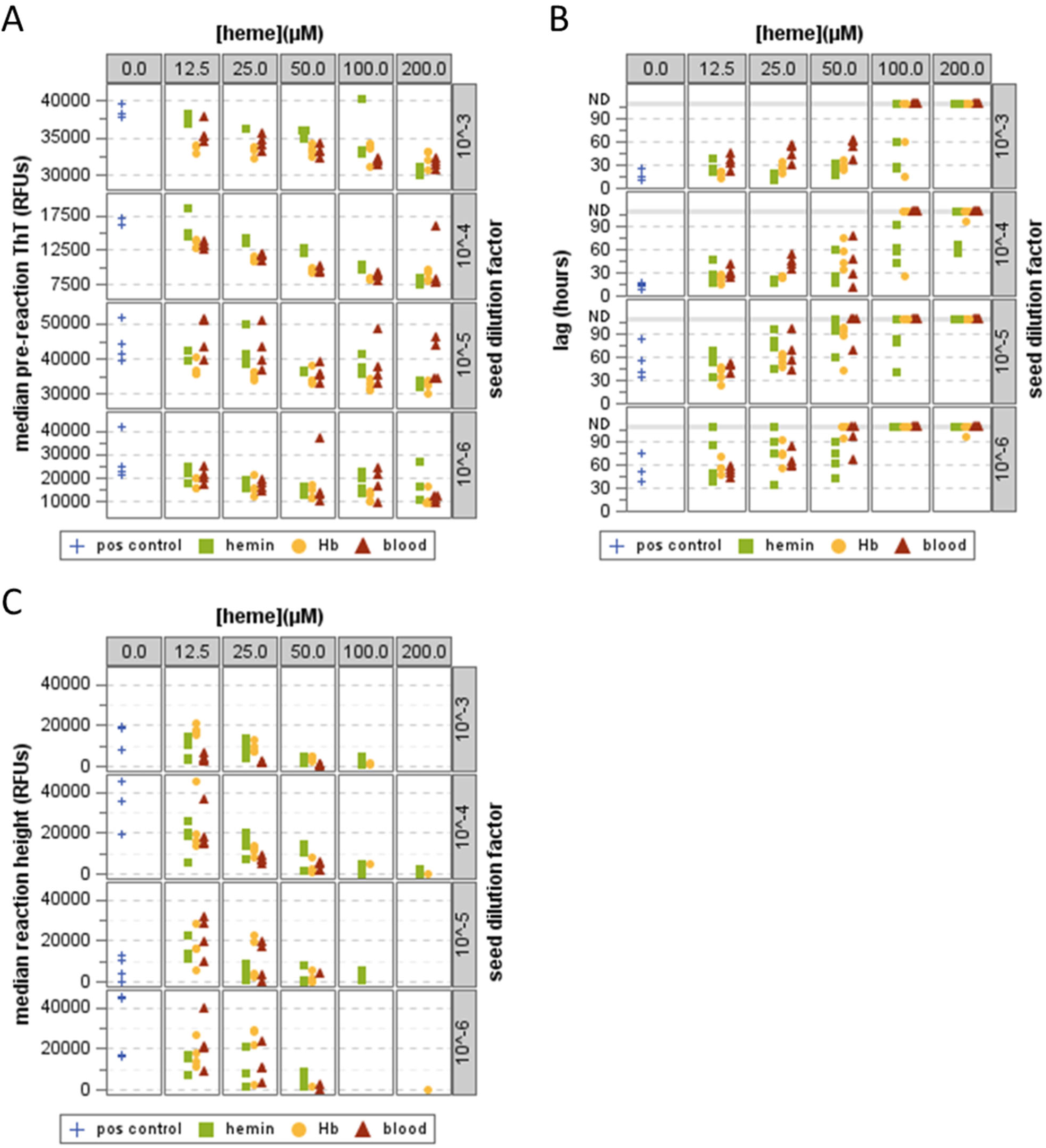
Effects of hemin, hemoglobin (Hb), and blood on RT-QuIC detection of Prions. Scatter plots depicting (A) pre-reaction ThT fluorescence, (B) reaction lag times, and (C) reaction ThT fluorescence. Reaction outcomes are shown at differing prion seed dilutions (rows), heme concentrations (columns), and heme inhibitor types (markers); with hemin shown as green squares, Hb as yellow circles, whole blood as red triangles, and buffer only controls as blue crosses. For reaction lag times (B) upper grey reference line denotes wells where no reaction was detected (ND).

The addition of heme, Hb, or blood to the seed homogenates resulted in dose-dependent increases in lag times at all seed dilutions (Fig 1b). At higher inhibitor concentrations, prion detection was completely ablated (Fig 1b). Of the three inhibitor sources tested, blood appeared to produce the greatest inhibition, followed by Hb, and then heme.

In addition to the longer lag times and lost detection of seeded-reactions, the morphology of the RT-QuIC reaction curves (ThT fluorescence vs. time) was changed by the addition of heme-containing inhibitors. In the presence of inhibitors, dose-dependent decreases were observed in the fluorescence signal, both during the pre-reaction period (Fig 1a) and following amplification (Fig 1c). Neither hemin nor Hb absorb photons strongly at the excitation or emission wavelengths of ThT (Hemin A_max_: 385 nm, Hb A_max_: 411 nm, ThT_ex_: 450 nm, ThT_em_: 480 nm), and others have demonstrated that the fluorescence of unbound ThT is not impacted by the presence of heme (25). To examine this more closely in our specific reaction conditions, we added the heme-containing inhibitors to RT-QuIC reaction buffer both with and without a standard amount (0.1 mg/ml) of rPrP^C^. Interestingly, while ThT fluorescence is known to increase sharply when bound to amyloid (36, 37), we did observe increased baseline fluorescence in the presence of rPrP^C^ (Fig 2a). The addition of heme-containing inhibitors showed minimal impact on ThT fluorescence in the rPrP^C^-free buffers but did suppress the rPrP^C^-dependent ThT fluorescence increase (Fig 2b). One interpretation of these results is that while hemin or Hb do not substantially interact with ThT alone, both molecules appear to interact with rPrP^C^. This interaction may be competitive, i.e., displacing ThT from rPrP^C^, or they may bind concurrently. In the latter scenario, the close proximity of the two bound molecules could allow the heme macrocycle to act as an acceptor for energy transfer, thereby suppressing ThT fluorescence(38).

**Figure 2.**
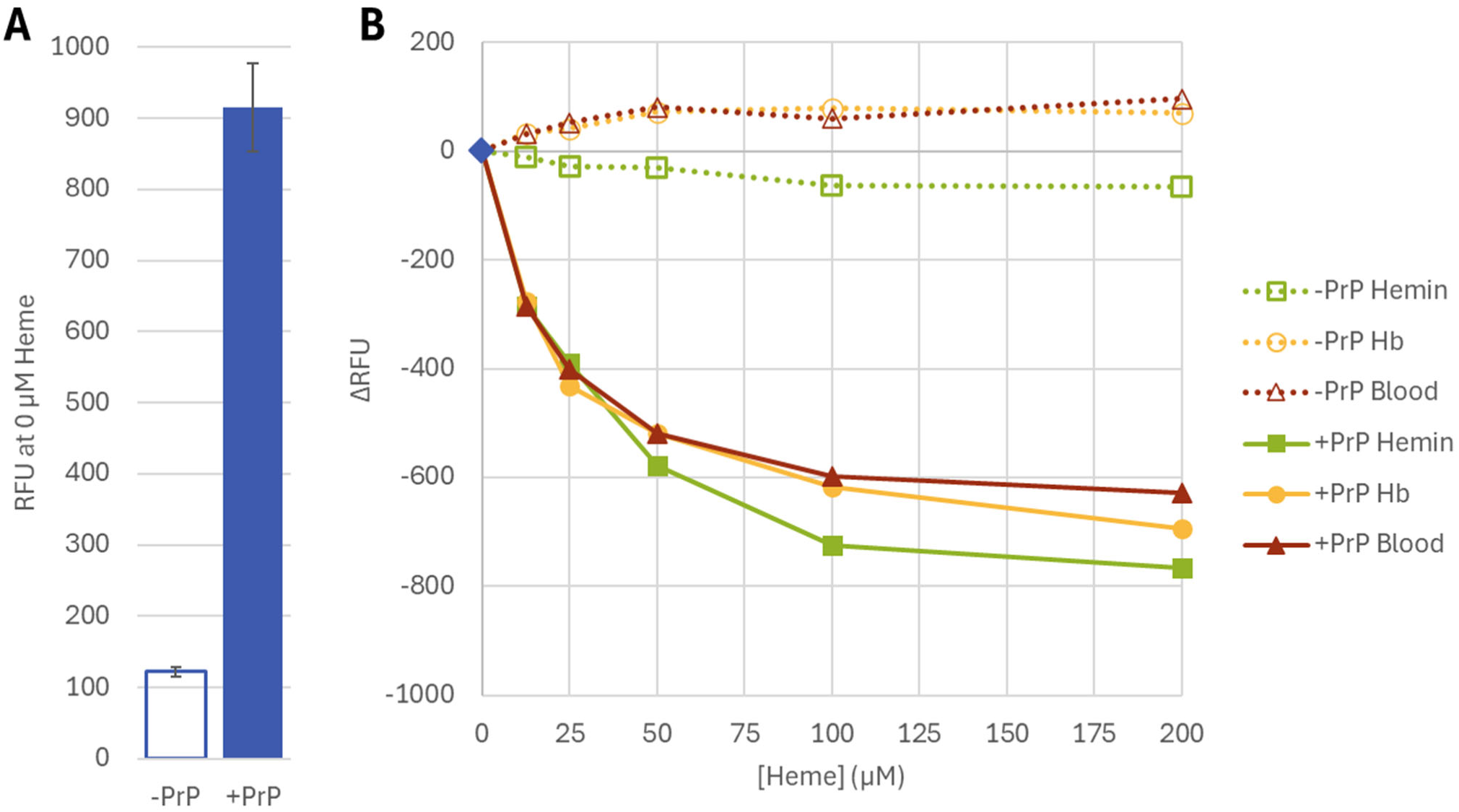
ThT interactions with rPrP^C^ and Heme-Containing inhibitors. Bar chart (A) displays mean fluorescence (RFU) for ThT alone, or in the presence of rPrP^C^. Error bars denote standard deviation (n=3). Line graph (B) shows change in ThT fluorescence (ΔRFU) measured in the presence of 12.5-200 µM hemin, Hb, or whole blood; with or without 0.1 mg/ml rPrP^C^. Values for hemin are denoted by green squares, Hb by yellow circles, and whole blood by red triangles. Mixtures containing rPrP^C^ are shown by filled markers and solid lines and reactions without by empty markers and dashed lines.

### Blood/Heme does not destroy PrP^Sc^ seeding activity

Previous research has demonstrated that exposure to heme results in the destruction or restructuring of other pathogenic amyloids (25–28). To test this potential with prions from a natural host, homogenate of a scrapie positive sheep brain was mixed 1:1 with either lysed whole blood, concentration-matched Hb solution, or 1× PBS and incubated at 4 °C for 1 week. These conditions were chosen to mimic those of a severely hemolyzed blood sample or heavily blood-contaminated tissue with subsequent refrigerated storage delays (24 h or 1 week) prior to analysis. The limiting dilution for detection was then determined using serial dilutions of each seed material. Except for the 10^−3^ dilutions, in which Hb was still present at >100 µM in the reaction mixture, significant differences in lag time or limits of detection were not observed (Fig 3). These results demonstrate that exposure to whole blood or hemoglobin in the tissue homogenate does not disrupt the seeding activity of PrP^D^ in the assay.

**Figure 3.**
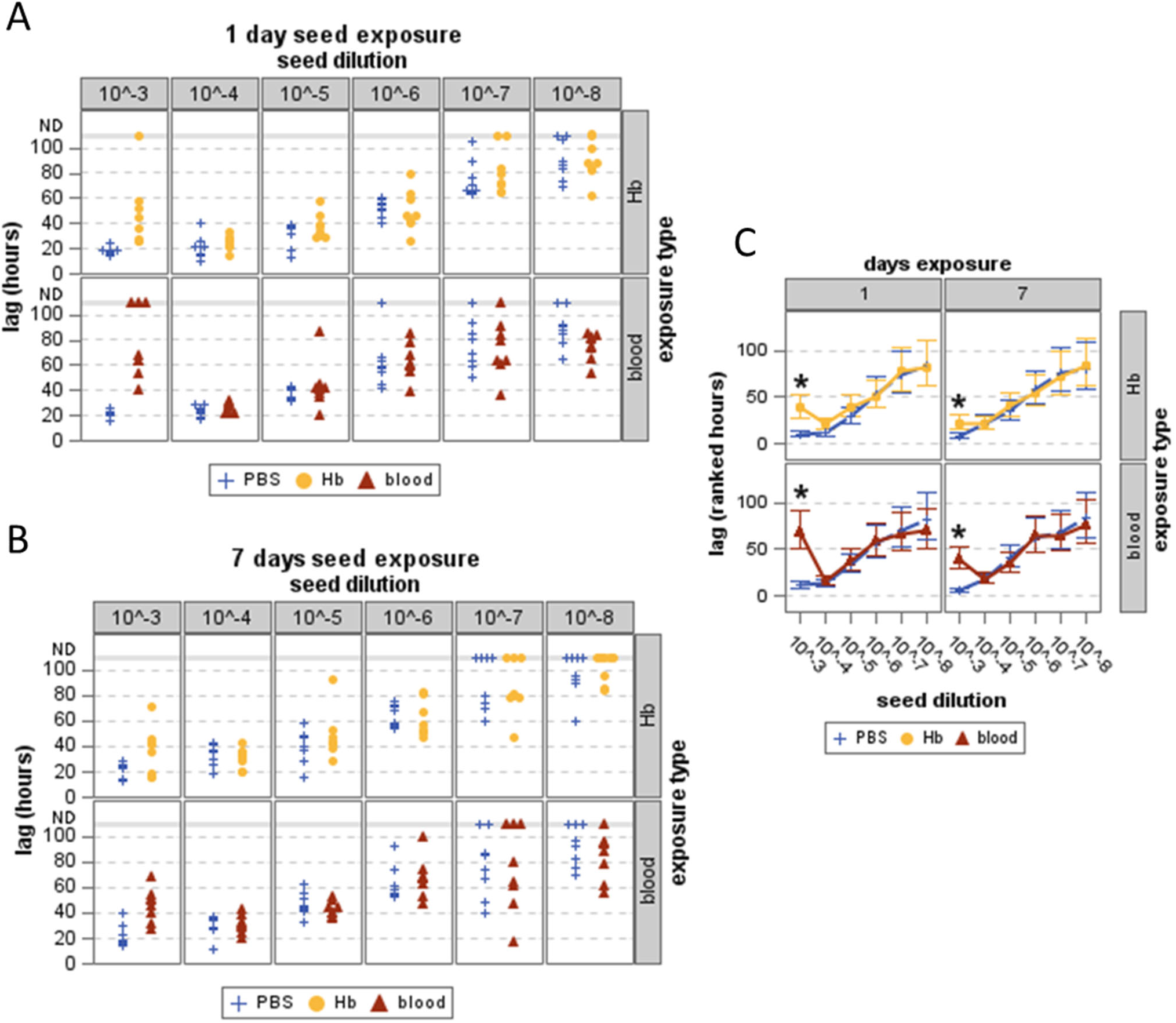
Seeding activity of PrP^Sc^ following exposure to Hb or Whole blood. Scatter Plots depicting reaction lag times for RT-QuIC following exposure of seed material to hemoglobin (Hb) (yellow circles) or whole blood (red triangles) for 1 day (A) or 7 days (B). Controls in each assay were exposed to PBS (blue crosses). Ranked lag times are also plotted against seed dilution factor (C) for each condition and its PBS control. Asterisks denote significant differences between ranked lag times from equivalent dilutions of inhibitor-exposed and PBS-exposed samples (all *P_Holm_* ≤ 0.0006).

### Inhibitors act on the rPrP^c^ assay substrate

Another fundamental component of amplification assays is the recombinant prion protein substrate. Prior research has shown that rPrP binds to heme in an isoform specific manner, where the heme:PrP binding ratio is higher for rPrP^C^ than for the misfolded form(23). To confirm similar heme:rPrP^C^ binding under RT-QuIC conditions, we spiked RT-QuIC reaction buffer (−ThT) [10 mM NaPO4 pH 7.4, 300 mM NaCl, 1 mM EDTA, 0.1 mg/ml rPrP^C^], with mock seed samples containing from 0-200 µM hemin. Binding of rPrP^C^ and hemin was observed as a red-shift in the maximum absorbance peak of heme (Fig 4a, b, d). In addition to the spectral shift, visible turbidity was evident in the spiked samples; this is observed as an overall increase in the UV-vis spectra absorbance baseline (Fig 4 a, b). Interestingly, significant turbidity was also observed in the 0 heme condition, where the reaction buffer was spiked with buffer [1× PBS + 0.1% SDS] alone (Fig 4a, c). To prevent interference from SDS-induced turbidity, heme spiking was also performed using a mock sample buffer of 1× PBS lacking SDS. These conditions did eliminate the turbidity observed at 0 µM hemin (Fig 4b,c) while maintaining the heme-binding-induced red-shift (Fig 4b). For this reason, the following experiments examining the binding ratio of hemin:rPrP^C^ were performed in the absence of SDS.

**Figure 4.**
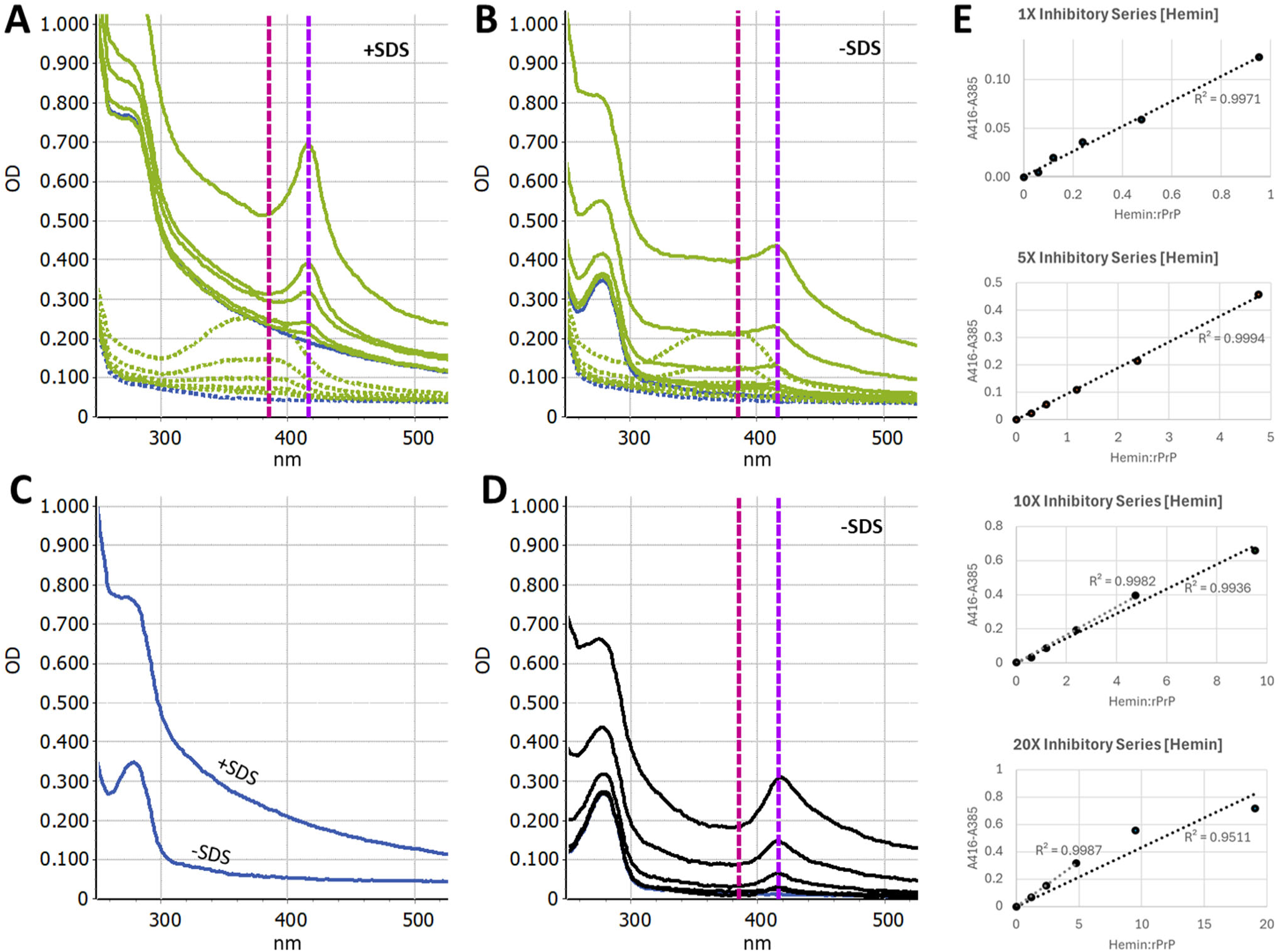
Ratio of Hemin:rPrP Interaction in RT-QuIC Buffer Conditions. (A,B) UV-Vis absorbance spectra of 0.1 mg/ml rPrP^C^ (solid blue), sample buffer only (dashed blue), hemin at 12.5, 25, 50, 100, and 200 µM (dashed green), and 0.1 mg/ml rPrP^C^ + Hemin [12.5-200 µM] (solid green); shown with SDS (A) and without SDS (B). (C) Spectra of rPrP^C^ and 0 µM hemin, with and without SDS. (D) Differential spectra of 0.1 mg/ml rPrP^C^ + Hemin [12.5-200 µM] with hemin spectra subtracted. The differential minimum (385 nm) and maximum (416 nm) are shown with dashed vertical lines (E) Hemin:rPrP^C^ binding ratios were tested with 0.1 mg/ml rPrP^C^ and multiples of the 12.5-200 µM inhibitory series hemin concentrations. The difference between the differential maximum and minimum (A416-A385) is plotted against the molar ratio of Hemin:rPrP^C^. Linear fits and accompanying R^2^ values are shown for whole (black dashed line) or partial (grey dashed line) series.

To quantify the spectral shift, differential spectra were calculated where the spectra of the hemin solutions alone was subtracted from those of the hemin+rPrP^C^ mixtures (Fig 4d). Evolution of a differential peak at 416 nm and a valley at 385 were observed (Fig 4d). The difference of these absorbance values was then plotted against the heme:rPrP^C^ molar ratio (Fig 4e). When rPrP^C^ was exposed to hemin at the previously tested inhibitory concentrations of 12.5-200 µM, a dose-dependent spectral shift was observed, however saturation was not reached. Subsequently, hemin was added at 5X, 10X, and 20X the inhibitory concentration series (i.e. 62.5-1000 µM, 125-2000 µM, and 150-4000 µM) to reach saturation. Ultimately, extensive precipitation of rPrP precluded precise quantification of the binding ratio; however, loss of linearity in the red-shift increase was seen beginning at molar ratios greater than ~5:1 hemin:rPrP^C^ (Fig 4e). This broadly agrees with the previously reported 7:1 binding ratio for heme to rPrP^C^ (23).

RT-QuIC reaction buffer (−ThT) was also spiked with Hb and whole blood at 12.5-200 µM. While binding of hemin to rPrP^C^ displays a strong shift in the differential (heme-subtracted) UV-Vis spectrum prior to incubation (Fig 5b), initial exposure of rPrP^C^ to Hb or whole blood results in a much smaller initial spectral shift (Fig 5c,d). This suggests the majority of heme in these conditions remains bound to Hb or that any transfer of heme to rPrP^C^ may occur more gradually over the assay run time. An example figure comparing the differential spectra to the raw and inhibitor-only spectra at the 200 µM conditions can be found in the supporting information (Fig. S1). To examine the inhibitor impacts at later assay timepoints under reaction conditions, RT-QuIC reaction buffer was spiked with sample buffer (SDS+) containing hemin, Hb, or whole blood and incubated at 42 °C for 24 h. These mixtures were then centrifuged at 21000 × g for 10 min to remove insoluble material and the resulting supernatants were analyzed by SDS-PAGE and Coomassie staining. In agreement with the initial spectroscopic measurements, even in the absence of heme, a significant portion of the rPrP^C^ substrate is lost after 24 hours at reaction conditions (Fig 5a). For the hemin-exposed samples, additional, dose-dependent rPrP^C^ loss was observed (Fig 5b). After 24 h, the Hb or blood spiked buffers unexpectedly showed more rPrP^C^ remaining in the buffer than in the 0 heme condition (Fig 5c,d), indicating that the presence of Hb or blood in the reaction mixture actually stabilizes the solubility of rPrP^C^. These results suggest that both hemin and Hb interact with rPrP^C^ in the assay substrate and impact its solubility in the reaction buffer, albeit through different mechanisms.

**Figure 5.**
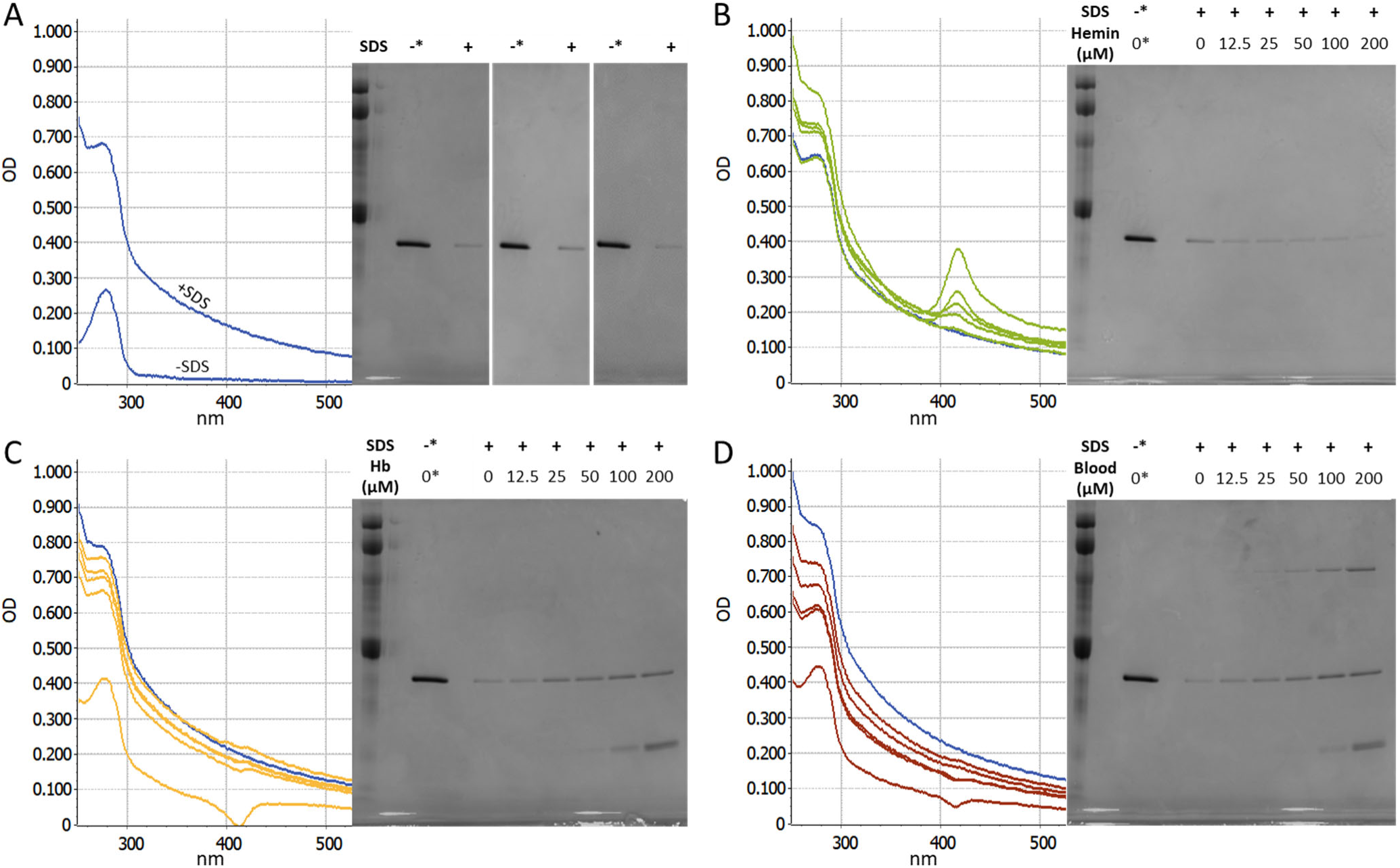
rPrP^C^ Substrate Stability in the Prescence of SDS and Heme-Containing Inhibitors. Differential UV-Vis absorbance spectra resulting from 0.1 mg/ml rPrP^C^ in RT-QuIC buffer (−ThT) exposed to mock seed material containing SDS (0.1% in sample buffer) (A) and Hemin (B), Hb (C), or Blood (D) at 0, 12.5, 25, 50, 100, and 200 µM. In panels B-D, the 0 µM condition is shown in blue. Also shown are SDS-PAGE and Coomassie staining of equivalent samples following incubation at 42°C for 24 hr and subsequent centrifugation to remove insoluble material. For Coomassie staining, control samples of 0.1 mg/ml rPrP^C^ in RT-QuIC buffer were prepared fresh and not incubated, denoted by asterisk.

In response to the evident changes to rPrP^C^ substrate stability following exposure to heme, heme-free RT-QuIC reactions were prepared using a range of substrate concentrations from 0.01 to 0.10 mg/ml (10-100%) so that the impact of rPrP^C^ depletion alone could be assessed. Reactions performed with depleted rPrP^C^, particularly those below 0.05 mg/ml (50%), exhibited longer lag times, lost detections, and diminished fluorescence signals (Fig 6) similar to the heme-mediated effects on the standard reaction mixture containing 0.1 mg /ml rPrP^C^.

**Figure 6.**
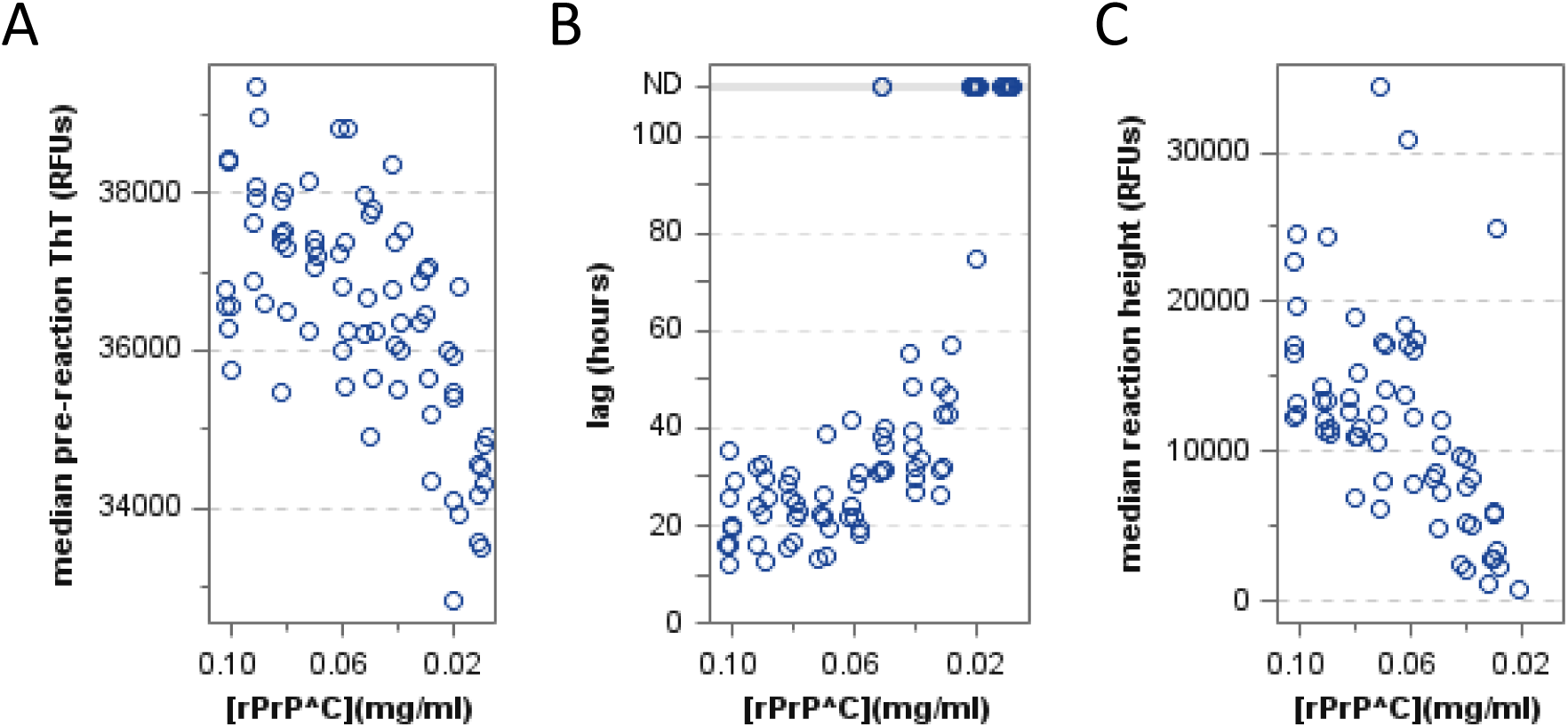
Effects of reduced substrate concentration on RT-QuIC detection of prions. Scatter plots depicting (A) pre-reaction ThT fluorescence, (B) reaction lag times, and (C) reaction ThT fluorescence at reduced rPrP^C^ substrate concentrations. Reactions were seeded at a 10^−4^ seed dilution in the absence of heme-containing inhibitors. For reaction lag times (B), upper grey reference line denotes wells where no reaction was detected (ND).

Lastly, a reaction condition matrix was prepared testing reactions containing 0.10, 0.15, and 0.20 mg/ml (100, 150, 200 %) rPrP^C^ spiked with 0, 50, 100, and 200 µM Hb. As readily observed in Figure 7A, reactions with elevated rPrP^C^ concentrations showed dose dependent rescue of lag times (Fig 7B), reaction frequency, and fluorescence intensity (Fig 7C), albeit with some delay in lag times in the absence of Hb (Fig 7C, 0 µM Hb). Together, these data demonstrate that interactions with the rPrP^C^ substrate are the primary factor by which heme-containing inhibitors disrupt prion detection by RT-QuIC.

**Figure 7.**
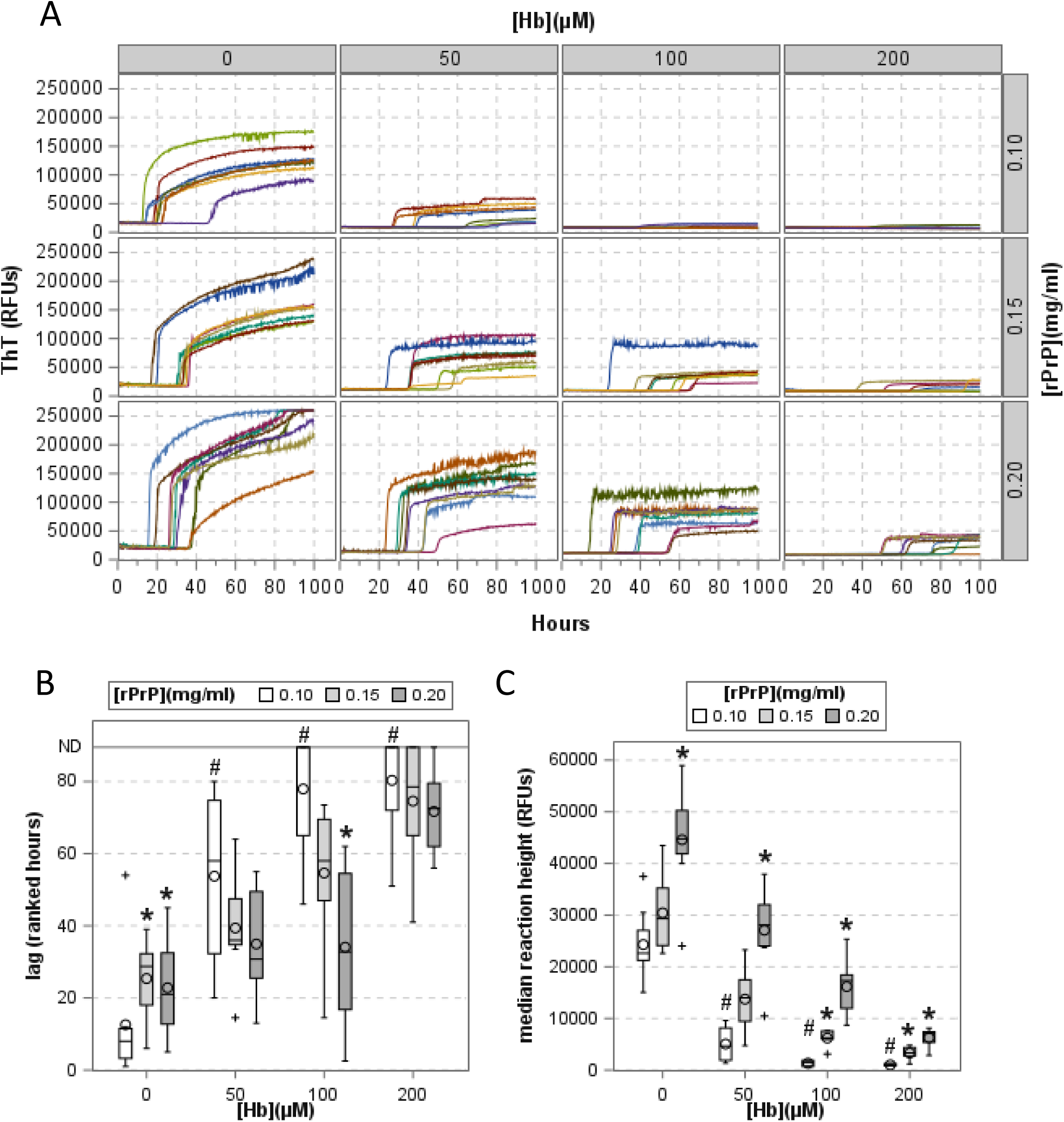
Rescue of RT-QuIC assay inhibition with by increased rPrP^C^ substrate concentration. (A) Raw ThT fluorescence curves for reactions seeded at a 10^−4^ seed dilution in the presence of varying Hb (columns) and increased rPrP^C^ substrate (rows) concentrations. Ranked lag times(B) and reaction ThT fluorescence (C) are shown as bar graphs where # denotes significant differences between Hb-containing and Hb-free reactions at 0.1 mg/ml rPrP^C^ (all *P_Holm_*< 0.0001) and * denotes significant differences between elevated (0.15, 0.20 mg/ml) and standard (0.1 mg/ml) rPrP^C^ concentrations within a given Hb concentration (all *P_Holm_* ≤ 0.005).

### Heme and protein constituents of Hb each contribute to inhibition of RT-QuIC

Given the apparent mechanistic differences between free hemin- and Hb-mediated inhibition, assays were also performed in the presence of apohemoglobin (apoHb) to extricate the respective contributions of the bound heme and globin protein components of Hb. ApoHb was prepared from the holoprotein (Hb) by acidified acetone extraction. Following re-constitution, apoHb concentrations were estimated using the previously published extinction coefficient ε_280nm_ = 0.0162 M^−1^ (29, 31). As apoHb is inherently unstable due to the lack of its native cofactor, absorbance-based protein quantification can be inexact. As a method of secondary confirmation, solutions containing equal amounts of Hb and apoHb (as estimated by 280 nm absorbance) were prepared and analyzed by SDS-PAGE and Coomassie staining. This comparison demonstrated grossly equivalent concentrations between the two forms (Fig 8a). Though the addition of apoHb did result in assay inhibition relative to 0 µM controls (Fig 8b), the overall effectiveness in prolonging lag time was generally less for the apoprotein (apoHb) as compared to the holoprotein (Hb) (main effect of Hb type: at 10^−4^, *P* < 0.0001; at 10^−5^, *P* = 0.0316). This was most evident when the reaction was seeded with 10-fold greater PrP^D^, where significant differences between Hb types were detected at 12.5 - 50 µM Hb (seeded at 10^−4^, *P_Holm_*≤ 0.0034), but only at 12.5 µM Hb when seeded at a dilution of 10^−5^ (*P_Holm_* ≤ 0.0211). These data suggest that both the globin protein component as well as the bound heme cofactor contribute to the inhibitory effects of Hb on RT-QuIC.

**Figure 8.**
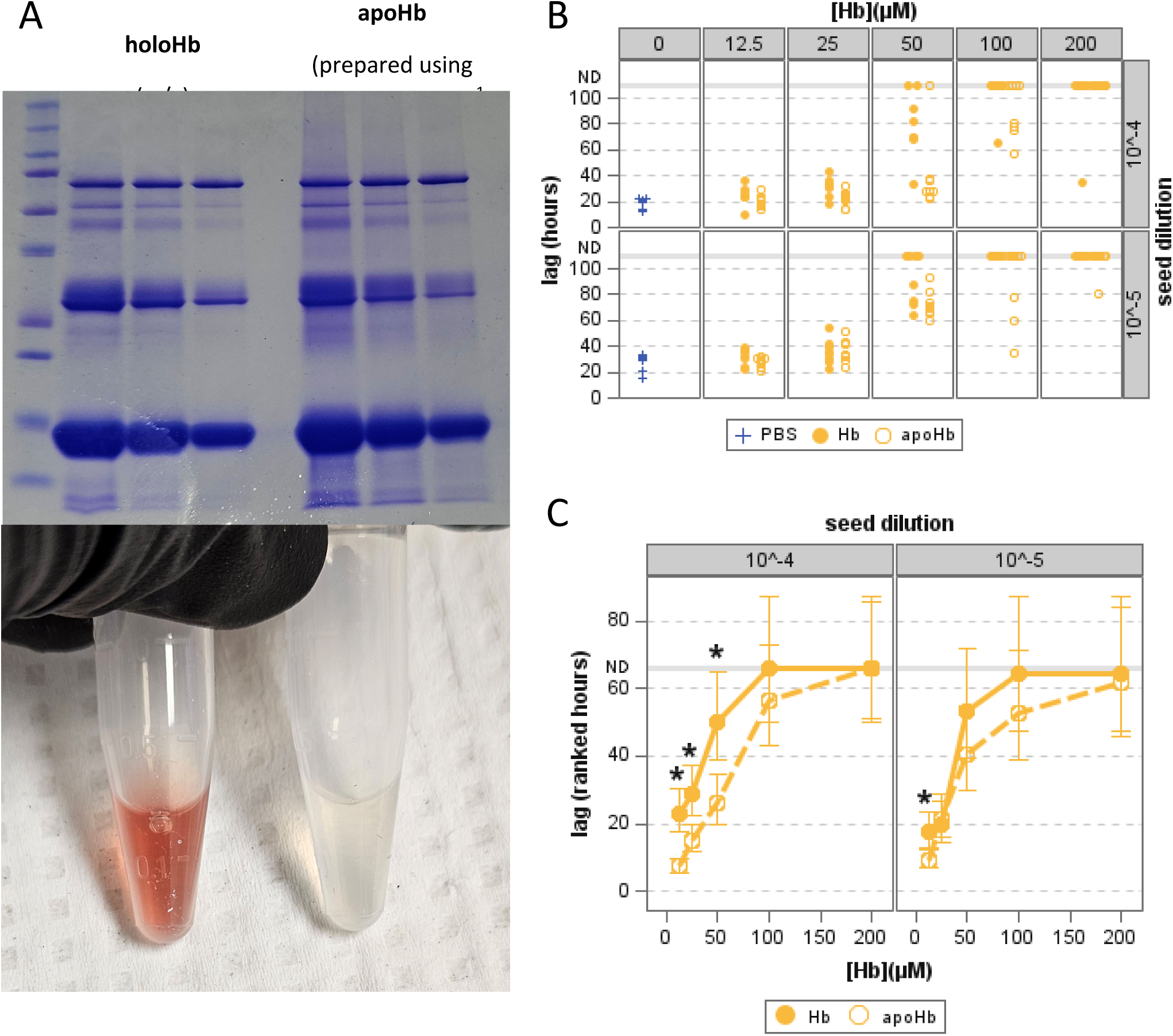
Comparative inhibitory effects of holohemoglobin(Hb) and apohemoglobin(apoHb). (A) Coomassie stained SDS-PAGE showing 1:2 dilution series of holoHb (left) and apoHb (right). (B) Scatterplot depicting reaction lag times of RT-QuIC reactions at differing seed dilutions (rows), Hb types (markers), and inhibitor concentrations (columns). (C) Ranked lag times plotted against inhibitor concentration. Filled yellow circles with solid lines show Hb and open yellow circles with dashed lines show apoHb. Asterisks denote significant differences between reactions containing equivalent concentrations of Hb or apoHb (all *P_Holm_* ≤ 0.034).

### Heme quantification in Diagnostic Tissues

To gauge the potential impact of heme/Hb-mediated inhibition on diagnostic efforts using RT-QuIC, various tissues from small ruminants were assayed for total heme concentration by oxalic acid fluorescence assay. The results of these measurements are summarized in Table 1, along with calculated extrapolations for various tissue homogenate dilutions. Briefly, while heme concentrations exceed inhibitory levels in nearly all intact tissues, by the time a 10^−3^ dilution was reached, only the blood, spleen, and placenta approached inhibitory levels of heme. However, several tissues maintained micromolar levels of heme if only diluted 10^−2^, which may still be capable of disrupting detection in peripheral tissues or during the earliest stage of infection where tissue samples are more likely to bear low prion titers.

**Table 1.** Heme Concentrations and Estimated Inhibition in Small Ruminant Tissues and Homogenate Dilutions. Heme concentrations measured by oxalic acid fluorescence assay. Values for homogenate dilutions directly measured in the assay are underlined. Heme concentrations shown for other homogenate dilutions and intact tissues are calculated from the bolded values. Some measured values fell outside of the standard curve linear range of 0-10 µM and are shown with a strikethrough. For placentas, five biopsies from each placenta were measured at a 1:500 dilution; mean values for each placenta are shown. Relative tissue homogenate concentrations are highlighted with a yellow color scale. Estimated heme-mediated inhibition levels are highlighted by a red color scale as shown at the bottom.

| Sheep #4645 |  |  |  |  |  | Goat #5010 |  |  |  |  |  |
| --- | --- | --- | --- | --- | --- | --- | --- | --- | --- | --- | --- |
| µM Heme in Tissues |  | Dilution relative to intact tissue |  |  |  |  | µM Heme in Tissues |  | Dilution relative to intact tissue |  |  |
| Tissue type | intact | 10<sup>-1</sup> | 10<sup>-2</sup> | 10<sup>-3</sup> | 10<sup>-4</sup> | Tissue type | intact | 10<sup>-1</sup> | 10<sup>-2</sup> | 10<sup>-3</sup> | 10<sup>-4</sup> |
| cerebellum | 216.35 | 21.63 | **<u>2.16</u>** | <u>0.12</u> | 0.02 | cerebellum | 190.15 | 19.02 | **<u>1.90</u>** | <u>0.11</u> | 0.02 |
| tonsil | 69.22 | 6.92 | **<u>0.69</u>** | <u>0.04</u> | 0.01 | tonsil | 255.62 | 25.56 | **<u>2.56</u>** | <u>0.14</u> | 0.03 |
| RPLN | 297.48 | 29.75 | **<u>2.97</u>** | <u>0.20</u> | 0.03 | RPLN | 297.86 | 29.79 | **<u>2.98</u>** | <u>0.16</u> | 0.03 |
| spleen | 15289.09 | 1528.91 | **<u>152.89</u>** | ~~14.79~~ | **<u>1.53</u>** | spleen | 14189.08 | 1418.91 | **<u>141.89</u>** | ~~15.61~~ | **<u>1.42</u>** |
| kidney | 355.43 | 35.54 | **<u>3.55</u>** | <u>0.26</u> | 0.04 | kidney | 905.46 | 90.55 | ~~11.66~~ | **<u>0.91</u>** | 0.09 |
| liver | 417.14 | 41.71 | **<u>4.17</u>** | <u>0.28</u> | 0.04 | liver | 899.72 | 89.97 | **<u>9.00</u>** | <u>0.79</u> | 0.09 |
| Brainstem | 98.34 | 9.83 | **<u>0.98</u>** | <u>0.07</u> | 0.01 | Brainstem | 77.29 | 7.73 | **<u>0.77</u>** | <u>0.05</u> | 0.01 |
| rectal mucosa | 60.89 | 6.09 | **<u>0.61</u>** | <u>0.05</u> | 0.01 | rectal mucosa | 6.76 | 0.68 | **<u>0.07</u>** | <u>0.00</u> | 0.00 |
| diaphragm | 373.90 | 37.39 | **<u>3.74</u>** | <u>0.24</u> | 0.04 | diaphragm | 983.58 | 98.36 | **<u>9.84</u>** | <u>0.62</u> | 0.10 |
| skeletal muscle | 487.72 | 48.77 | **<u>4.88</u>** | <u>0.55</u> | 0.05 | skeletal muscle | 943.83 | 94.38 | ~~12.35~~ | **<u>0.94</u>** | 0.09 |
| CSF (supernatant) | 6.32 | **<u>0.63</u>** | <u>0.04</u> | 0.01 | 0.00 | CSF (supernatant) | -0.04 | 0.00 | **<u>0.00</u>** | <u>0.01</u> | 0.00 |
| Blood | 8758.43 | 875.84 | 87.58 | **<u>8.76</u>** | <u>0.58</u> | Blood | 7868.67 | 786.87 | 78.69 | **<u>7.87</u>** | <u>0.43</u> |

| Sheep #4649 |  |  |  |  |  | Sheep Placenta |  |  |  |  |  |  |
| --- | --- | --- | --- | --- | --- | --- | --- | --- | --- | --- | --- | --- |
| µM Heme in Tissues |  | Dilution relative to intact tissue |  |  |  |  | µM Heme in Tissues |  | Dilution relative to intact tissue |  |  |  |
| Tissue type | intact | 10<sup>-1</sup> | 10<sup>-2</sup> | 10<sup>-3</sup> | 10<sup>-4</sup> | placenta number | intact | 10<sup>-1</sup> | 10<sup>-2</sup> | 1:500* | 10<sup>-3</sup> | 10<sup>-4</sup> |
| cerebellum | 20.78 | 2.08 | **<u>0.21</u>** | <u>0.00</u> | 0.00 | #1266 | 2266.27 | 226.63 | 22.66 | **4.53** | 2.27 | 0.23 |
| tonsil | 932.53 | 93.25 | **<u>9.33</u>** | <u>0.75</u> | 0.09 | #1267 | 1645.78 | 164.58 | 16.46 | **3.29** | 1.65 | 0.16 |
| RPLN | 1785.91 | 178.59 | ~~17.46~~ | **<u>1.79</u>** | 0.18 | #1268 | 1732.77 | 173.28 | 17.33 | **3.47** | 1.73 | 0.17 |
| spleen | 14824.96 | 1482.50 | **148.25** | ~~17.40~~ | **<u>1.48</u>** | #1269 | 1340.98 | 134.10 | 13.41 | **2.68** | 1.34 | 0.13 |
| kidney | 382.37 | 38.24 | **<u>3.82</u>** | <u>0.23</u> | 0.04 | #1270 | 2071.36 | 207.14 | 20.71 | **4.14** | 2.07 | 0.21 |
| liver | 639.23 | 63.92 | **<u>6.39</u>** | <u>0.42</u> | 0.06 | #1271 | 2748.37 | 274.84 | 27.48 | **5.50** | 2.75 | 0.27 |
| Brainstem | 32.85 | 3.29 | **<u>0.33</u>** | <u>0.01</u> | 0.00 | #1272 | 2318.99 | 231.90 | 23.19 | **4.64** | 2.32 | 0.23 |
| rectal mucosa | 56.51 | 5.65 | **<u>0.57</u>** | <u>0.01</u> | 0.01 | #1273 | 2085.75 | 208.58 | 20.86 | **4.17** | 2.09 | 0.21 |
| diaphragm | 416.14 | 41.61 | **<u>4.16</u>** | <u>0.25</u> | 0.04 | #1274 | 2565.11 | 256.51 | 25.65 | **5.13** | 2.57 | 0.26 |
| skeletal muscle | 452.87 | 45.29 | **<u>4.53</u>** | <u>0.27</u> | 0.05 | #1275 | 1322.32 | 132.23 | 13.22 | **2.64** | 1.32 | 0.13 |
| CSF (supernatant) | 0.57 | **<u>0.06</u>** | 0.00 | 0.00 | 0.00 | Average | 2009.77 | 200.98 | 20.10 | 4.02 | 2.01 | 0.20 |
| Blood | 8738.70 | 873.87 | 87.39 | **<u>8.74</u>** | <u>0.53</u> |  |  |  |  |  |  |  |

**Table 1. Heme Concentrations and Estimated Inhibition in Small Ruminant Tissues and Homogenate Dilutions.**
| Heme inhibition (µM) |  |  |
| --- | --- | --- |
| 0 |  |  |
| 12.5 |  | <u>Underlined</u> = directly measured |
| 25 |  | **Bold** = used to calculate |
| 50 |  | ~~Strikethrough~~ = measured but outside linear range |
| 100 |  |  |
| 200 |  |  |

## Conclusions

In this study we demonstrate that each of heme, Hb, and whole blood inhibit the RT-QuIC assay. Delayed lag times were observed with seed material samples containing as low as 12.5 µM heme and, in the case of lower prion-titer samples, a substantial loss of replicate detections was observed at concentrations as low as 50 µM for Hb or whole blood. Heme itself, the globin component of Hb, and the remaining constituents of whole blood are each contributing factors to the observed inhibition. When prion containing samples were incubated in the presence of Hb or whole blood prior to being assayed, seeding activity was not lost, suggesting that inhibition occurs at the level of the assay itself. Hemin was confirmed to be able to bind rPrP^C^ in RT-QuIC reaction buffer, and both free hemin and Hb induced solubility changes in the protein substrate. When heme-free reactions were performed with limited quantities of substrate, outcomes mirror those of reactions inhibited by heme, Hb, or blood. Finally, increasing the starting rPrP^C^ concentration was able to rescue Hb inhibited prion detection, reinforcing the conclusion that inhibition from Hb or blood containing samples is a result of depletion of available rPrP^C^ substrate in the assay mixture.

When heme was measured in small ruminant tissues, levels were, not unexpectedly, shown to vary between tissue types. Of interest, nearly all types of intact tissue contain >200 µM heme; however, given our data suggesting pre-reaction exposure does not degrade PrP^D^ seeding activity, the concentrations at the actual testing dilutions are more relevant. At the most commonly tested tissue dilution, 10^−3^, only the blood, spleen, and placenta still contained millimolar quantities, and even these tissues would be at the low end of the inhibitory range. It should be noted that the tissues in this study were collected from freshly euthanized animals which were then exsanguinated and necropsied in a timely manner. It is therefore possible that field collected samples may have a considerably greater degree of blood contamination or hemolysis. Even so, given the results from whole blood and splenic tissue, it remains unlikely that heme-mediated inhibition would be more than a minimal factor at a 10^−3^ dilution or greater. While these levels of sample dilution are common practice, they are inherently limiting to the theoretical sensitivity of the assay. In recent years, a variety of efforts have been made to enrich or purify prions from tissues (10, 39–41), some of which are only semi-selective for PrP^D^ and may result in increased inhibitor concentrations along with the enriched prions. Other strategies aim to allow for testing of less diluted samples (42, 43). In these cases, it is possible that heme or Hb levels may again be present at inhibitory levels. Somewhat conveniently, the ability to see the color of a solution of aqueous Hb with the naked eye roughly coincides with the inhibitory ranges described in this study (Fig 9). As a broad observation, if the final seed material to be introduced to the RT-QuIC assay is visibly red/pink/brown colored, Hb-mediated inhibition may influence the reaction lag times, fluorescence signal maxima, and frequency of detection, and thus should be considered when interpreting results.

**Figure 9.**
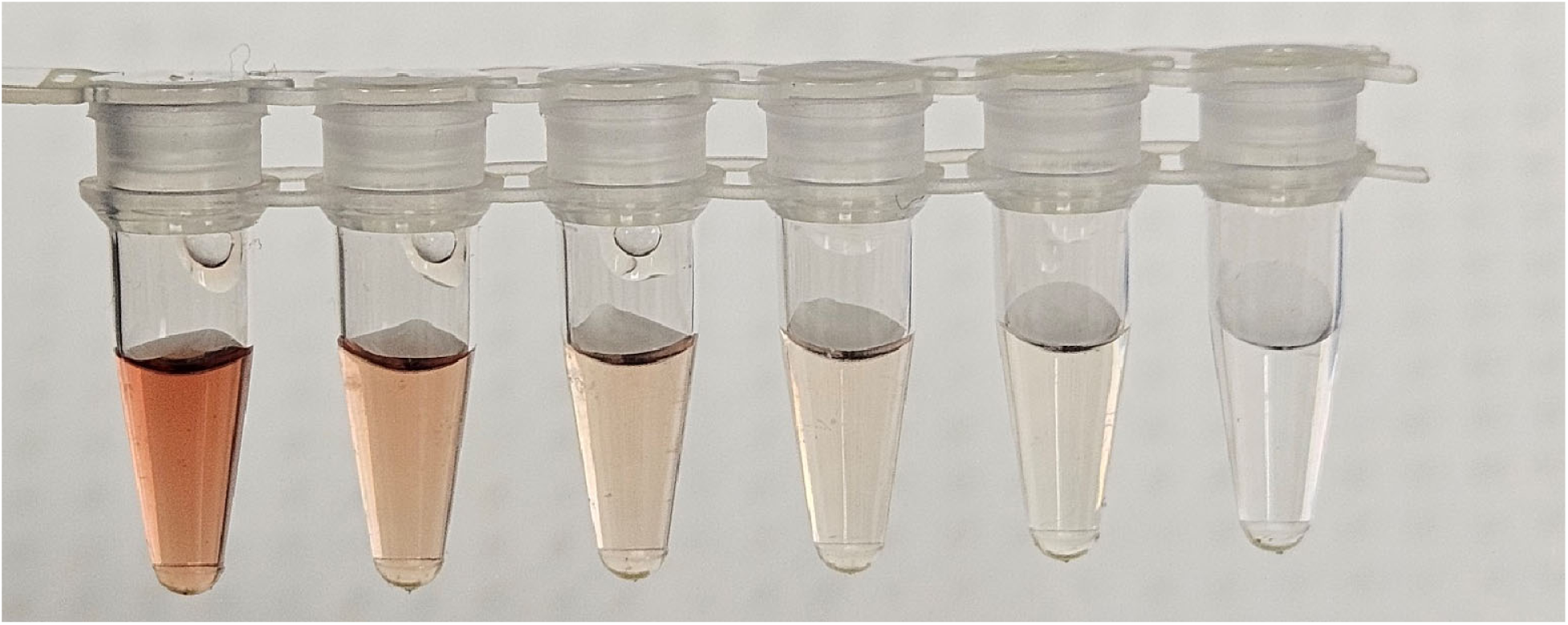
Visual appearance of inhibitory Hb concentrations. Hb in Sample Buffer at 200, 100, 50, 25, 12.5, and 0 µM.

## Acknowledgements

The authors wish to acknowledge Lori Fuller and the animal care staff at the Animal Disease Research Unit and Washington State University and thank them for their excellent technical and animal care support. The research presented in this manuscript was funded by USDA-ARS Project Number 2090-32000-042-000-D, https://www.ars.usda.gov/research/project/?accnNo=441236. Mention of trade names or commercial products in this article is solely for the purpose of providing specific information and does not imply recommendation or endorsement by the US Department of Agriculture.

---

**Figure S1.**
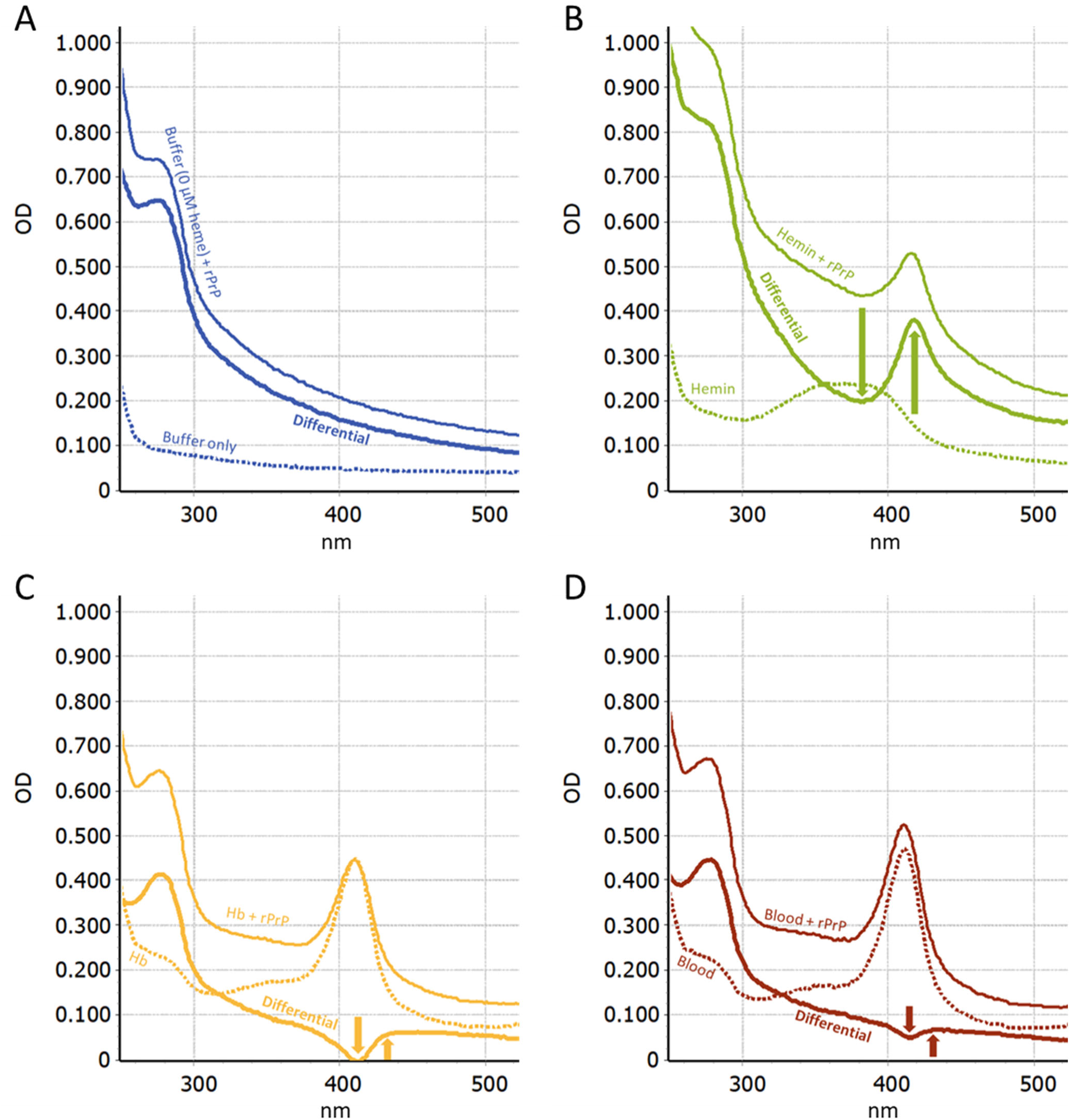
Differential spectra and spectral shifts resulting from inhibitor interactions with rPrP^C^ in RT-QuIC Buffer. UV-Vis absorbance spectra of 0.1 mg/ml rPrP^C^ exposed to various sources of heme at 200 µM; sample buffer (0 µM heme, SDS+) only (A), Hemin (B), Hb (C), or Blood (D). Each panel shows the raw absorbance spectrum for the heme source alone (dashed), the raw absorbance spectrum for the heme-spiked rPrP^C^ (solid), and a differential spectrum where the heme source spectrum is subtracted from the mixture spectrum (bold). Shifts in the differential spectra are highlighted with arrows.

